# From qualitative to quantitative insect metabarcoding: an in tandem multilocus mosquito identification methodology

**DOI:** 10.1101/2020.11.22.393140

**Authors:** Katerina Kassela, Adamantia Kouvela, Michael de Courcy Williams, Konstantinos Konstantinidis, Maria Goreti Rosa Freitas, Andreas Nearchou, Elisavet Gatzidou, Stavroula Veletza, Georgios C. Boulougouris, Nikolaos Dovrolis, Ioannis Karakasiliotis

## Abstract

In the era of emergence and re-emergence of vector-borne diseases, a high throughput trap-based insect monitoring is essential for the identification of invasive species, study of mosquito populations and risk assessment of disease outbreaks. Insect DNA metabarcoding technology has emerged as a highly promising methodology for unbiased and large-scale surveillance. Despite significant attempts to introduce DNA metabarcoding in mosquito or other insect surveillance qualitative and quantitative metabarcoding remains a challenge. In the present study, we have developed a methodology of in-tandem identification and quantification using cytochrome oxidase subunit I (COI) combined with a secondary multilocus identification and quantification involving three loci of 28S ribosomal DNA. The presented methodology was able to identify individual species in pools of mosquitoes with 95.94% accuracy and resolve with high accuracy (*p* = 1, *χ*^*2*^ = 2.55) mosquito population composition providing a technology capable of revolutionizing mosquito surveillance through metabarcoding. The methodology, given the respective dataset, has the potential to be applied to various small animal populations.

## Introduction

Vector borne diseases constitute a public health concern worldwide, accounting for more than 17% of all infectious diseases^1^. Among the most important infectious disease vectors are mosquito species of the genera *Anopheles*, transmitting malaria, *Culex*, transmitting West Nile virus and *Aedes*, transmitting Dengue, Yellow fever, Chinkungunya and Zika virus^2, 3^. Not all mosquito species transmit pathogens or at the same rate. Authorities worldwide organize entomological surveillance activities in order to identify changes in insect population dynamics and the emergence of invasive species. Morphology based taxonomy is the most common approach used in species identification; however, it has important limitations. The greatest challenge is the discrimination of morphologically similar species or specimens with damaged external features. Besides, the accuracy of the morphological approach is based on the level of expertise and is time consuming^4^.

DNA barcoding is a taxonomic molecular method, usually implicating the cytochrome oxidase subunit I (COI) gene for animal species discrimination^4^. In cases where COI is not sufficient, other taxonomically important genomic loci are used as 12S mtDNA, 16S mtDNA, Cytochrome B ITS, 18S and 28S rDNA, EF-1a and NADH^5-10^. DNA metabarcoding is a recently developed molecular approach that engages next-generation sequencing (NGS) into a high-throughput identification of species in a mixed population^11^. DNA metabarcoding has already been applied in many different taxonomic groups, such as plants and aquatic organisms, fungi, insects, and mammals^12^ ^13^ ^14^. A recent review summarizes the importance, requirements and challenges for a successful metabarcoding approach for insects, envisioning an automated smart-trap that may simultaneously collect and identify insect populations through coupled in-trap metabarcoding^15^. COI metabarcoding methodologies have inherent limitations towards a quantitative approach as the COI gene shows extensive third-base drift even within the same species^16, 17^. Mismatches may result in decreased PCR amplification efficiency that leads to an unpredictable taxon/species amplification bias and skewing of relative representation of a certain taxon during quantification^14, 17, 18^. Multiplexing regions of COI during metabarcoding offered an approach that aimed to smooth out amplification bias with an identification efficiency of ∼80%^19, 20^. An alternative approach using 28S ribosomal RNA was able to distinguish mosquitoes more efficiently^21^. However, methodologies developed up to date are far from efficient in quantitative metabarcoding.

While qualitative metabarcoding based on COI has been successful enough in species identification, a quantitative metabarcoding approach remains the Holy Grail for the efficient insect population surveillance. In the present study, we developed a novel methodology for qualitative and quantitative analysis of mosquito populations through in-tandem multilocus metabarcoding.

## Results and Discussion

A quantitative metabarcoding procedure requires a sequence database of all species in an area. In order to construct a database of ribosomal RNAs we performed total RNA sequencing (RNA-seq) on morphologically identified specimens assisted by single-specimen COI DNA barcoding. Twenty-four different mosquito species were sequenced, most of which for the first time (**Suppl. Table 1**). Ribosomal RNAs and COI were assembled and used to populate the rRNA/rDNA and COI reference databases (**Suppl. Table 1**). The databases were further expanded to include *Aedes echinus* and *Anopheles plumbeus* that were later analysed by Sanger sequencing. Multiple alignment of all reference rRNAs resulted in a map of conserved and hypervariable regions (**Figure 1**). 18S rRNA was highly conserved amongst species and was not used further in the study (**Figure 1**). On the other hand, 28S rRNA revealed isles of conserved and hypervariable regions (D2, D8 and D10 in *Diptera*) (**Figure 1**) known before for their ability to distinguish closely related species. The D2 region has been used in the taxonomy of *Acarina, Hymenoptera, Heteroptera* and *Diptera*, including mosquitoes^21-27^. D8 and D10 regions have been used to a lesser extent in the classification and phylogeny of mites and mealybugs^22, 27, 28^. Conserved regions flanking hypervariable regions were used for the design of three pairs of universal oligonucleotides (100% identity) (**Suppl. Table 2**). A pair of degenerate primers, targeting a 329 bp region of COI, was used to distinguish all mosquito species in the local and the BOLD database (**Suppl. Table 2**).

**Figure 1.**
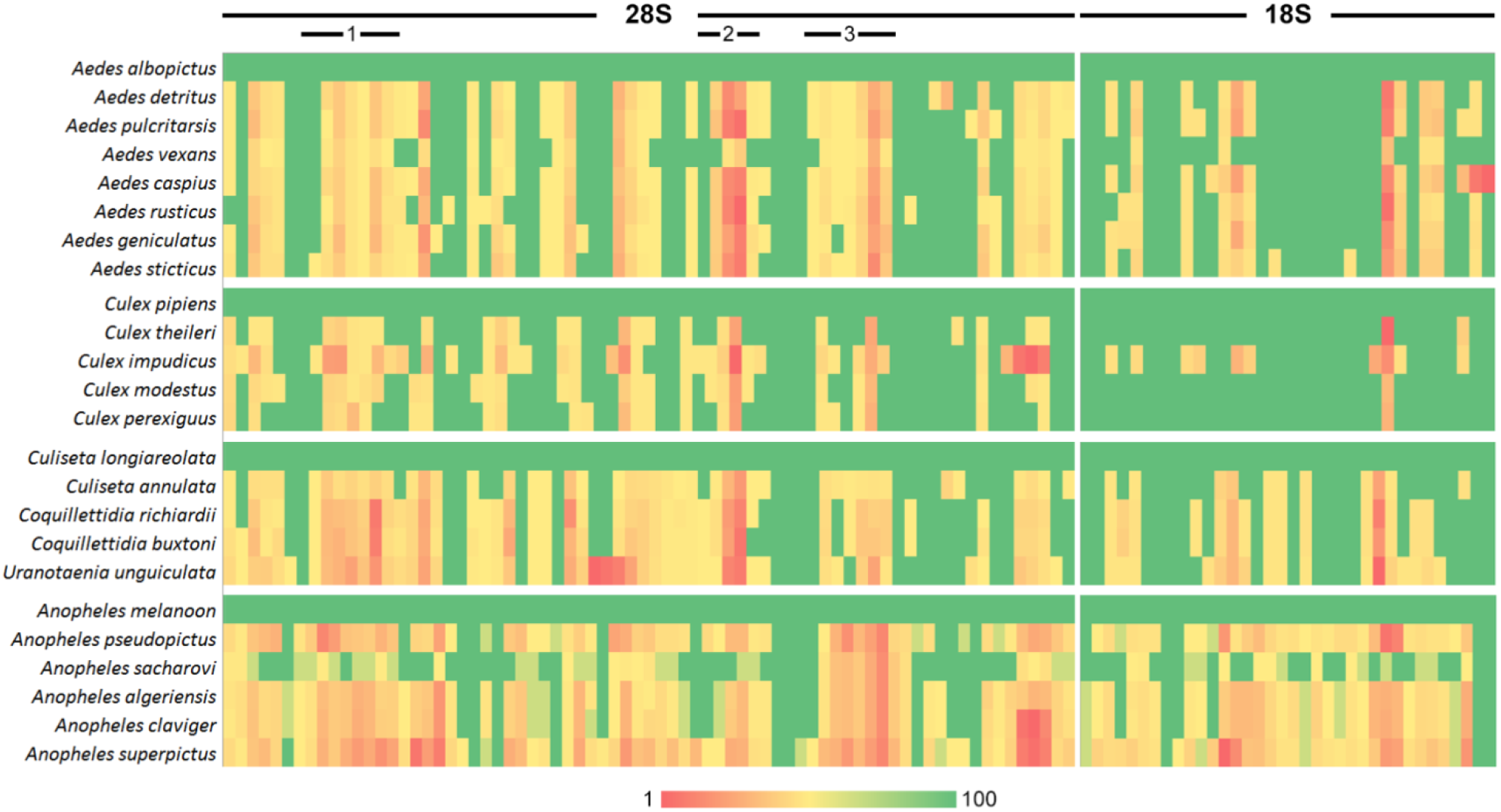
Similarity heatmap of 28S and 18S rRNAs coloured according to similarity percentage (1-100 %) per 60 bp of multiple alignment. The first species in each group was used as reference. Indicators (1,2 and 3) correspond to the three 28S regions used in the study (n=24 species).

As DNA content varies greatly among species, quantitative analysis requires a training set (custom pools of individuals with known population composition) for the development of a mathematical model. Twenty-seven pools composed of different proportions of 22 species were prepared (**Suppl. Table 3**). Four rare species that were not represented by enough individuals served as negative controls. Extracted DNA from the pools was subjected to PCR amplification with the set of the three 28S primer pairs one targeting COI. All four PCR products, corresponding to the same pool, were processed as a single NGS DNA library and sequenced using 400-bases chemistry. Small reads were a significant disadvantage of previous methodologies^19, 20^. Reads were filtered for truncated amplicons and mapped on the COI database. Reads mapping on a specific COI were used to calculate haplotype networks within a pool (**Figure 2**). Reads with significantly divergent haplotypes (<98%) were inspected as new haplotypes may reflect rare, invasive or cryptic species. Although it may not be possible to track back to an individual after metabarcoding, genetic similarity may be used as a guide in the subsequent identification of the new species with follow-up traps in the same habitat. For example, a species identified by morphological analysis (pool 22) as member of the *Anopheles maculipennis* group, appeared as a distant variant of *Anopheles melanoon* (**Figure 2C**).

**Figure 2.**
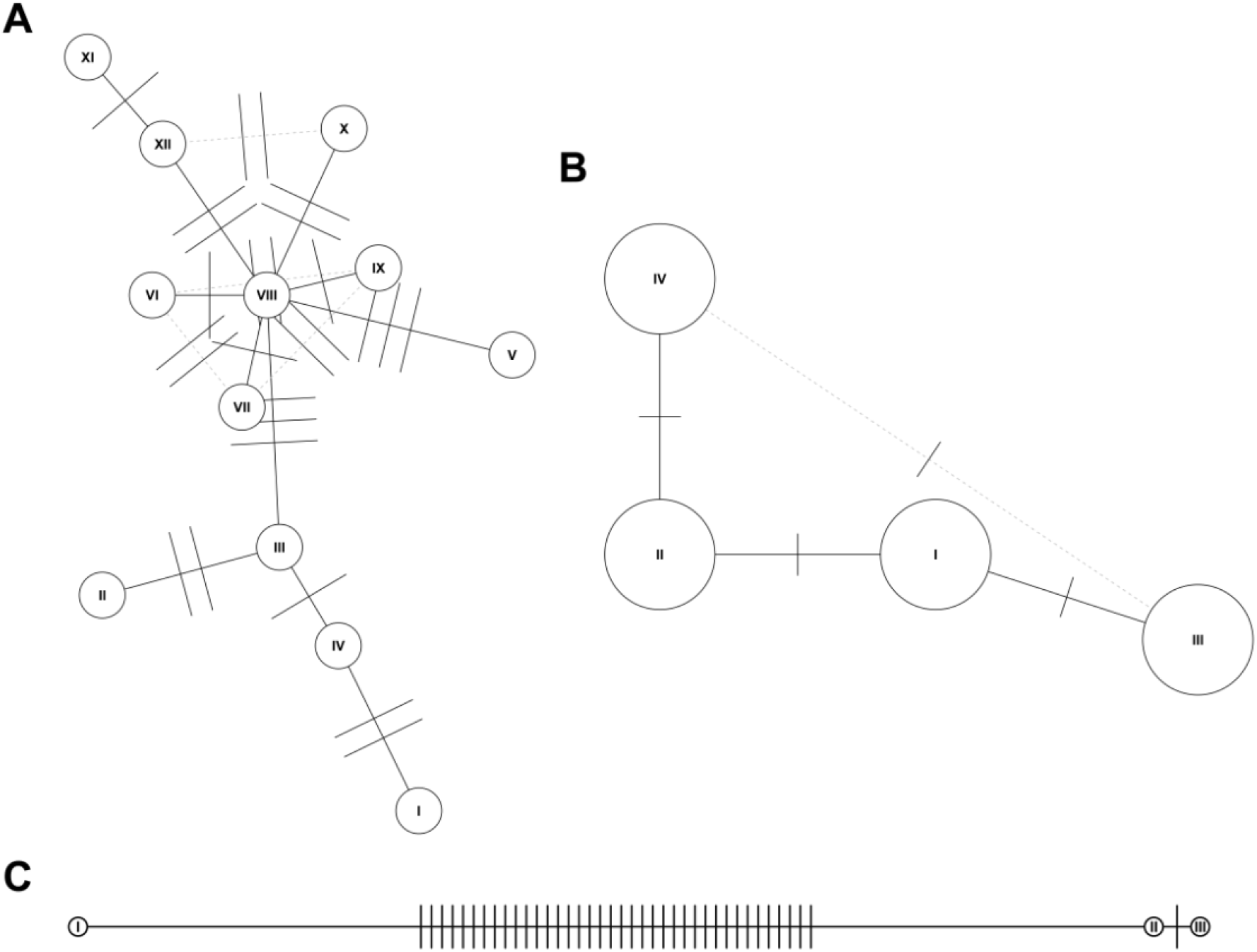
COI haplotype networks representing COI variability per species within a species [or allocated reads per species] in a pool of mosquitoes. (A) *Aedes caspius* haplotype network in pool 18 [n=79]. (B) *Aedes albopictus* haplotype network in pool 19 [n=12]. (C) *Anopheles melanoon* haplotype network in pool 22 [n=3].

Species identified through COI were used to build a temporary rDNA reference for each pool. Total reads were mapped on the three 28S rDNA loci within this rDNA temporary reference. Mapped reads for each of the three 28S rDNA loci were expressed as relative abundance reads per mosquito individual amongst species in a pool. The median relative abundance for all mosquito species was calculated after combining the results of all custom pools, yielding a mathematical constant (median read ratio) for each species per primer pair. The proposed model for the estimating mosquito abundance from the relative number of reads was based on a proportionality ansatz. This model of proportionality between reads and abundance corresponds to a set of linear equations for the relative abundance of species in a pool. The set consists of one equation per species, whereas the requirement for satisfying all equations was described in the form of a Matrix-vector representation 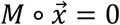 (**Eq. 1**).

*M* is a non-symmetric, square matrix with dimensionality equal to the number of species in a pool. The elements of *M* are functions of the number of reads, and the proportionality ratio between reads and number of individuals per species. For example, a pool with three different species and three mapped loci (A, B, C) Eq. 1 takes the form of Eq. 2.

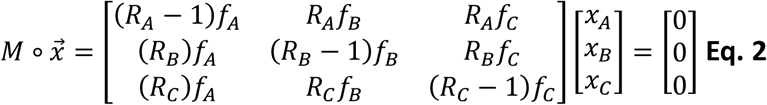

Where *R*_*A*_ = *r*_*A*_/Σ_*α*_ *r*_*α*_ is the ratio of the total reads of the mapped locusA (*r*_*A*_) to the total number of reads for all mapped loci of that pool, and *f*_*A*_ = *r*_*A*_/*N*_*A*_ is the ratio of total reads of the mapped locus A to the number of individuals of a species *N*_*A*_. The proposed approach is based on solving simultaneously the set of equations for each locus. An estimation of the number of mosquitoes for each species tagged with a specific locus is expressed in vector form as the solution, 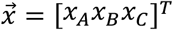 of Eq 2.

The calculated median read ratios were used to predict the composition of the pools (**Figure 3**). For the estimation of the population composition of each pool, the respective data were excluded from the training set. In order to measure the efficiency of our model we did a *χ*^*2*^ goodness of fit analysis on the predicted data used to create the histograms of mosquito abundance. In **Figure 3B, C** we present the *p* and *χ*^*2*^ values for the *χ*^*2*^ goodness of fit analysis for each of the pools examined. Note that the number of degrees of freedom in each experiment for the *χ*^*2*^ goodness of fit may vary between the different pools since it is equal to the number of species in the pool minus 1 (given that we do not impose any additional constrain in the moments of the expected discreet distribution of mosquito abundance). The three primer pairs demonstrated different efficiencies in predicting population composition. Primer pair 1 was the most efficient (median *p*= 0.789 and median *χ*^2^ =3.882) followed by primer pair 3 (median *p*= 0.743 and median *χ*^2^= 4.449) and primer pair 2 (median *p*= 0.464 and median *χ*^2^= 10.380) (**Figure 3B**,**C**). Combination of all three primer pairs demonstrated superior quantification efficiency (median *p* = 0.997, max =1, min = 0.196 and median χ^2^ = 1.147, min = 0.059, max = 11.588). In terms of identification efficiency, the pipeline resulted in a 95.94% accuracy, which is significantly higher than previous endeavours, attributed to the multiple correction checkpoints, while the overall statistical strength of the method reached *p* = 1 and *χ*^*2*^ = 2.55. To date, this methodology presents a significant advancement in qualitative mosquito identification in terms of accuracy while it is the first to assess unbiased insect quantification efficiently.

**Figure 3.**
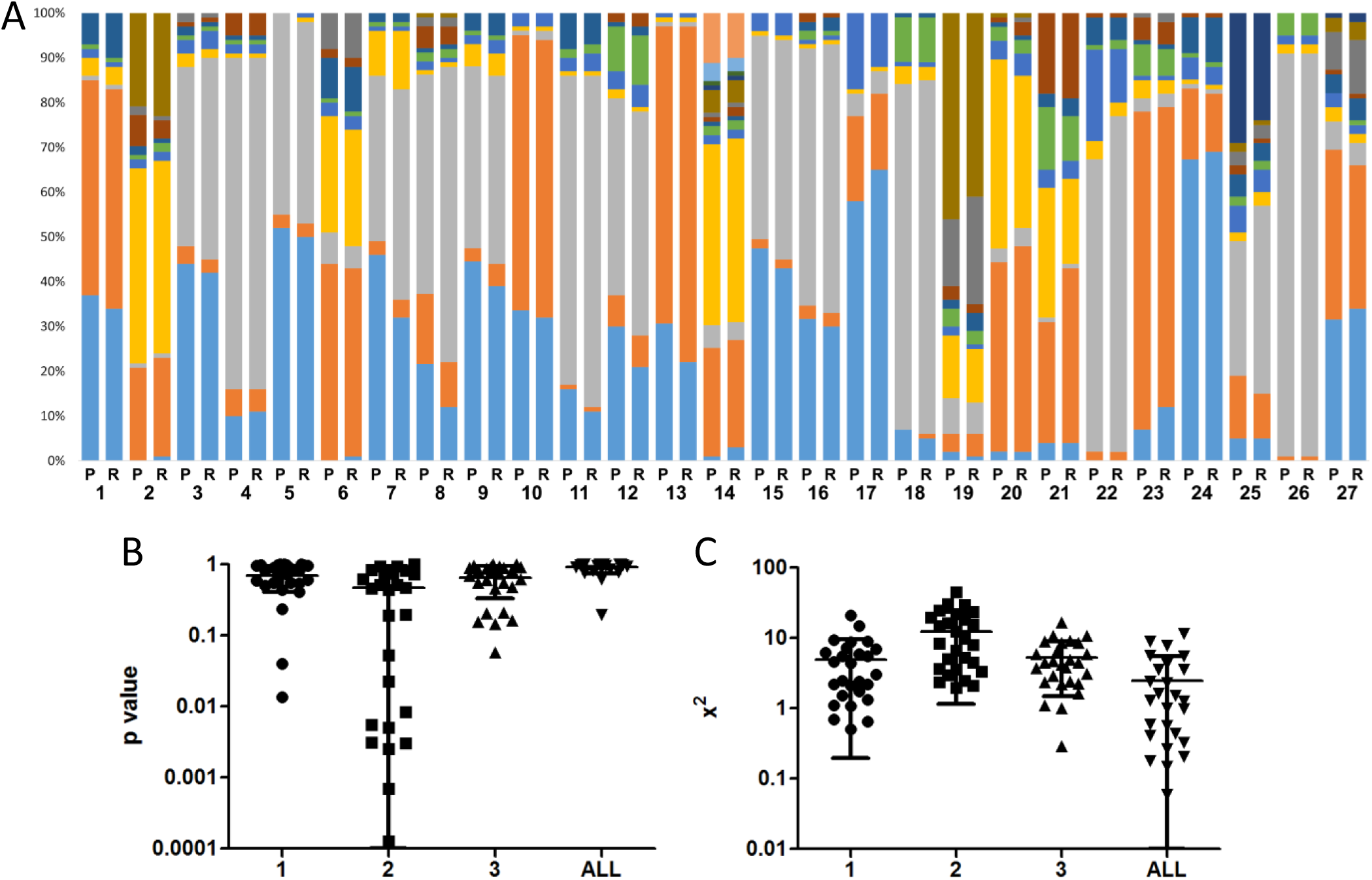
(A) Stack bars represent the diversity of each pool as predicted [P] by the metabarcoding pipeline compared to the actual [R] composition of the pool as determined through morphological and single-specimen COI barcoding. Numbers 1-27 represent the trap numbers while colour coding depicts species variation only between P and R of the same trap. (B) *P* value and (C) x^2^ vertical scatter plots of the three independent quantifications (1,2,3) based on the three 28S primer pairs and their combination (ALL) [n=27 pools, mean and standard deviation error bars].

## Conclusions

In tandem multilocus identification may constitute a potent methodology for qualitative and quantitate analysis of insect populations. Addressing a significant number of methodological issues and concerns described by previous reports^15^, such as long read length chemistry, proportional read representation and decoupling quantification from COI identification, we achieved high identification and quantification efficiency in pools of variable complexity. The methodology offers a platform globally adaptable when local or global training sets are provided, while it may serve as a guideline for similar metabarcoding approaches in other small animals. The same methodology is anticipated to complement and guide the identification of novel and cryptic species and subspecies through haplotype divergence networks. Overall, quantitative metabarcoding will scale up and revolutionize the way we perform insect surveillance and insect population genetics.

## Methods

### Mosquito collection and identification

Adult mosquitoes were collected using Centers for Disease Control (CDC) light traps with CO_2_. Mosquito specimens were examined over a bed of crushed ice at all times to maintain their condition, both during sample sorting and in making species identifications. Samples were otherwise stored at -80 °C prior to RNA and DNA extraction. Female mosquitoes were identified using external morphological features. Species nomenclature follows Harbach, 2018^29^ and generic abbreviations follow Wilkerson et al.^30^. Morphological identification was done using a combination of the keys^31, 32^ and the online resource MosKeyTool^33^. Representatives of the *Anopheles maculipennis* group cannot be distinguished morphologically among adult females^33^ and specimens were identified prior to use with COI barcoding from an excised leg.

### Species identification through COI barcoding

Where appropriate DNA barcoding was done using standard COI PCR and Sanger Sequencing. Mosquitoes were homogenized and total DNA was extracted by TRIzol reagent (Thermo Fischer Scientific) according to the manufacturer protocol. Universal primers COI_F and COI_R were used to amplify a 600 bp PCR product. The PCR reaction mixture contained 0.25x GC buffer, 1.5 mM MgCl_2_, 1 mM dNTPs mix, 0.2 μM of each primer and 1.5 U KAPA Taq DNA polymerase (Kapa Biosystems). The thermal profile of the PCR included 40 cycles of denaturation at 95°C for 30 s, annealing at 50 °C for 45 s and elongation at 65°C for 1 min, and a final elongation step at 65°C for 7 min. PCR products were purified using the NucleoSpin Gel and PCR Clean-up purification kit (Macherey-Nagel). Sanger Sequencing was performed on the PCR product and analyzed using the Barcode of Life Data System V4 platform ^35^.

### Total RNA Next Generation Sequencing

Mosquitoes were homogenized and total RNA was extracted by TRIzol reagent (Thermo Fischer Scientific) according to the manufacturer protocol. Whole transcriptome libraries were prepared from 500 ng of RNA extract, using the Ion Total RNA-Seq v2 Core Kit (#4479789, ThermoFisher Scientific) according to manufacturer instructions. In brief, the RNA library preparation involved RNA fragmentation, adapter ligation, reverse transcription and 14 cycles of PCR amplification using Ion Xpress™ RNA-Seq Barcode 1-16 Kit (#4475485, ThermoFisher Scientific). Quantification of the library was performed using Qubit Fluorometer high-sensitivity kit (ThermoFisher Scientific) and its median size was determined in LabChip GX Touch 24 (PerkinElmer). The libraries were loaded into an Ion 540 chip, using the automated Ion Chef System (Thermo Fisher Scientific) and sequencing was carried out on an Ion GeneStudio S5, ion torrent sequencer (ThermoFisher Scientific).

### Ribosomal RNA de novo assembly

Raw sequences from the pools of each of the mosquito species, with a member count of n=5 for each pool, were used as input for RNA-seq de novo assembly using Trinity v2.8.5 ^36^. Trinity, based on the de Bruijn graph algorithm^37^, produces contigs (set of overlapping DNA segments) that represent alternate transcripts of genes while treating sequences with structural changes (mutations and indels) as isoforms of the same gene. The whole process is performed via 3 distinct modules, namely Inchworm, Chrysalis and Butterfly, each respectively responsible for creating the assemblies of transcripts, clustering them and optimizing the de Bruijn graphs. These contigs were subsequently used as input to custom BLASTn^38^ queries to identify and annotate the transcripts that correspond to the 28S, 18S and 5.8S rRNA genes. Due to the absence of known rRNA sequences in Genbank^39^ for most of the mosquito species, contig length was used as an indicator of identity, based on already annotated mosquito species. To validate and refine these findings, the raw sequencer output was aligned on a custom reference created by the annotated rRNA contigs using the STAR tool^40^. This step produced alignment BAM files that after manipulation with samtools^41^ were converted to fastq files. These files underwent a second round of assembly, as previously described, allowing for more complete and accurate rRNA transcripts.

### Identification of conserved and hypervariable rRNA regions

All the assembled 28S and 18S rRNAs were aligned via ClustalW multiple sequence alignment^42^. Pairwise similarity per 60 bases (per ClustalW line) for all mosquito species was calculated using a custom Python script and plotted as a heatmap. Regions with absolute similarity amongst genera flanking hypervariable regions within a range of 450 bases were selected for the design of 3 sets of universal 28S rRNA oligonucleotides. 18S rRNAs showed high levels of conservation amongst related species and were not used further in the study.

### Mosquito pool preparation and DNA extraction

Pools of 100 morphologically identified mosquitoes (assisted by DNA COI barcoding) were prepared. Each pool consisted of various mosquito species at different proportions. The pools were homogenized in 3 ml of lysis buffer (50 mM Tris pH 8.0, 100 mM EDTA, 100 mM NaCl, 1% SDS). DNA was extracted from 1 ml of crude homogenate containing 6 μl of proteinase K (22 mg/ml, Macherey-Nagel), incubated overnight at 55°C.

### PCR amplification for DNA seq

Extracted DNA from each pool (50 ng) was used for the amplification of three distinct regions in the 28s rDNA and one region in the COI gene. The thermal profile of the PCR for the three 28s fragments included 20 cycles of denaturation at 95 °C for 30 s, annealing at 54 °C (28s mosq1)/56 °C (28s mosq2, mosq3) for 30 s and elongation at 72 °C for 30 s. A final elongation step was performed at 72°C for 5 min. The PCR program for COI amplification (COImosq F/R primers) included 35 cycles of denaturation at 95 °C for 30 s, annealing at 58 °C for 20 s and elongation at 74 °C for 30 s, and a final elongation step at 74°C for 5 min. The PCR reactions were carried out by KAPA Taq DNA Polymerase (KAPA biosystems). PCR amplification cycles were optimized in order to correspond to the logarithmic phase of each reaction. Equal quantities of the four PCR products from each mosquito pool were mixed and purified using Agencourt AMPure XP (Beckman Coulter).

### PCR product Next Generation Sequencing

Libraries were prepared from 100 ng of pooled amplicons, using the Ion Plus Fragment Library Kit (#4471252, ThermoFisher Scientific) according to manufacturer instructions, along with Ion Xpress™ Barcode Adapters kit (#4474518, ThermoFisher Scientific). Barcoded libraries were purified using Agencourt AMPure XP (Beckman Coulter) and quantified using Qubit Fluorometer high-sensitivity kit. Ion 530 chips were prepared using Ion Chef System (Thermo Fisher Scientific) and NGS reactions were run on an Ion GeneStudio S5, ion torrent sequencer (ThermoFisher Scientific).

### Quantitative species identification

To identify and quantify the mosquito species in mixed pools a two-pronged approach was employed involving sequencing of PCR products of COI and the three 28S rDNA regions. The sequencing reads that resulted from the above multiplex NGS were used for species identification using a bioinformatics pipeline of four distinct steps.

Step 1: Filtering of the sequences. A lower limit of 240bp length was imposed on the reads in order to avoid heavily truncated by-products. This limit was calculated based on the median read length of the samples and the genomic region of our primers. The filtering limit was implemented using the reformat script that is part of the BBMAP suite^43^ on each of the pool samples. This script does not trim the sequence string but rather selects sequences longer than the limit.

Step 2: Alignment to the 3’ COI database. In the local COI database, each locus represented the 3’ COI of the respective species. The bwa-mem v. 0.7.17-r1188 aligner^44^ was used to align all the filtered reads to the custom COI database discarding reads which map to multiple loci (secondary reads). In order to avoid sequencing errors and potential contaminations, we used a cut-off of less than half of the reads that represented an individual mosquito of the respective species. The output of this step was a list of species represented by adequate reads within a specific pool.

Step 3: Custom rDNA reference design. For each pool of mosquitoes, we created a custom reference file from our novel 28S rDNA database. Each reference file contained the three amplicon reference sequences for the identified-by-COI species. These custom reference files were constructed by parsing the output of the previous step of identified species by COI and retrieving the amplicon sequences matching the appropriate terms (species name) from our rDNA database. These actions were performed using bash scripts and commands.

Step 4: Alignment to the custom rDNA database. The algorithm, implemented as a new script, in this step first indexed (bwa index) the custom reference for each pool created in the previous step, then aligned (bwa-mem) the raw reads of the pools to the respective index (without secondary alignment) and finally retrieved the read counts for each species (for all 3 hypervariable regions) via samtools idxstats. These counts were piped into our mathematical model described below, to quantify the mosquitoes per species in our pools.

### Within-population diversity

A bioinformatics pipeline was constructed to identify population diversity within members of the same species using NGS reads, as an extension of the previous steps. When this pipeline was applied to both COI and rDNA reads it allowed us to visualize the genetic heterogeneity of individual mosquitoes. After the alignment of mosquito pool samples with our COI and rDNA databases it allowed us to isolate reads of a specific species and work just with those, refining the temporal and computational cost. The first step was a quality control/filtering algorithm that discards truncated reads and removes the read region where PCR primers bind in order to avoid false readings by their degeneration. This process was performed using a combination of samtools, BBMAP’s reformat and SeqKIt v. 1.2-r94 ^45^. These filtered reads were then clustered based on a 100% similarity threshold using CD-HIT v.4.8.1 ^46^. CD-HIT allows for a more manageable size of sequences to be used as input for our analyses and also helps identify sequences that are just products of sequencing errors (since the latter don’t represent a significant percent of similar sequences). By introducing a lower limit for member-sequences per cluster, we could identify only the clusters that contained well-represented sequence variations. The representative sequences of each filtered cluster were selected and used to create a custom .fasta file that undergoes multiple sequence alignment using MUSCLE v3.8.31 ^47^. The aligned .fasta file was used as input for the pegas package ^48^ in R for creating haplotype networks to visualize the differences and the evolutionary relationships among our sequences. Pegas creates networks using a randomized minimum spanning tree algorithm and provides an ability to visualize the sequence relationships, including their nucleotide differences, after clustering them into haplotypes.

### Mathematical model

Estimation of the relative abundance of mosquito species in a pool was based on a model of proportionality between the relative abundance of mosquito species in the pool and the expectation/average value of the number of mapped reads (filtered truncated amplicons) of a locus of a species in a pool where the same protocol has been used. This model of proportionality between reads and relative abundance corresponds to a set of linear equations for the relative abundance of mosquito species in a pool described in the form of a Matrix-vector representation in the form of Eq. 1. In order to drive the system of linear equations that show this relation for each of the species in a pool in the form of *Eq. 1* shown above we start by introducing our basic set of variables used to quantify the relationship. For any given pool we therefore represent *r*_*α*_ *as* the total reads of the mapped locus *α, and N*_*a*_ *as the* number of mosquitoes tagged based on locus α, extended over all loci where the number of reads exceeds the threshold. Based on those two experimentally measured variables we define as *f*_*a*_ = *r*_*a*_/*N*_*a*_ the ratio of total reads of the mapped loci *α* (*r*_*a*_) to the number of mosquito of the corresponding species *N*_*α*._ It is now possible to express the proposed proportionality relation of our model as relationships between our auxiliary variables of any two loci *A,B* in the form of *Eq. 3*

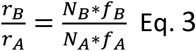

For clarity reasons in the final representation we also introduce as auxiliary variables, the ratio of reads for each tagged locus A as *R*_*A*_ = *r*_*A*_/Σ_*α*_ *r*_*α*_. We also introduce a set of reduced variables for both the total reads *r*_*α*_ and the *f*_*a*_ratios, by dividing them by the corresponding values of a reference locus that can be chosen arbitrary, resulting in a new set of reduced variables 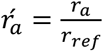 and 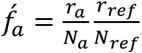, where the suffix *ref* corresponds to the locus that we have chosen as reference. It can be demonstrated that the reduced ratios are identical to the original ratios *R*_*a*_ = *Ŕ*_*a*_∀*a* and most importantly that the formal solution of Eq. 1 is not effected by the choice of the reference. The final system of linear equations comes from the estimation of the expected value for each of the ratios *Ŕ*_*A*_ in the form of Eq. 4.

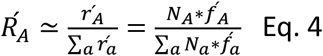

Since for each pool there will be one equation of the form of Eq.4 for each tagged locus we are able to form the system of linear equations that connects *Ŕ*_*A*_ and *Nα* in the form of *Eq. 1* and *Eq. 2* described above.

The core of the proposed model lies on the estimation of expected (i.e. the average) values for reads to mosquito ratio ⟨*f*_*a*_⟩ from pools with known mosquito abundance. Once the elements of the matrix *M* are estimated based on the expected values of the proportionality ratio between reads and the mosquito number *f*_*a*_ ≃ ⟨*f*_*a*_⟩ is known for the given experimental setup we can predict the unknown mosquitoes abundance for each species by, solving Eq 1. It turns out that the solution of *Eq 1* in terms of linear algebra is possible by estimating the vectors that when multiplied by the matrix *M* will result in a zero vector (i.e. will belong in what is called the null space of Matrix *M*). The elements of this vector 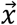 are estimate for the relative mosquito abundance. If we know the total number of mosquitos in a given pool, we can further impose the condition that relative mosquito abundance must be rational (ratio of integers). In addition, the elements of the vector 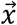 must be integers and sum up to the total number of mosquitos in the pool. To find the optimal solution that satisfies this restriction we start from our unconstrained estimate of vector 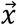 and find the mosquito abundance ratio that minimizes the square error of *Eq 1:* 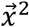. Therefore, the overall scheme can be understood as a list square fit of the total number of reads and the mosquito abundance in the proposed model, where the total number of reads are expected to be proportional to the abundance of each species.

Note that for the resulting system of equations, the formal solution of the problem is a vector in the null space of a matrix and therefore is not affected by the multiplication of a row by a constant and this means that the use of a reference tagged locus does not affect the formal solution. In practice, we use a numerical implementation of the singular value decomposition of the resulting matrices to estimate an initial mosquito abundance ratio, followed by the minimization of error, under the condition that the number of mosquitoes have to be integers as described above.

## Acknowledgments

This research has been co-financed by the European Union and Greek national funds through the Operational Program Competitiveness, Entrepreneurship and Innovation, under the call RESEARCH – CREATE – INNOVATE (project code:T1EDK-5000).

## Supplementary Tables

**Supplementary Table 1.** GenBank accession number table for 28S, 18S rRNA and COI complete or partial sequences

| Species | GenBank accession number |  |  |
| --- | --- | --- | --- |
|  | 28S rRNA | 18S rRNA | COI |
| *Aedes albopictus* | MT808421 | MT808449 | MT993476 |
| *Aedes caspius* | MT808431 | MT808459 | MT993477 |
| *Aedes detritus* | MT808422 | MT808450 | MT993478 |
| *Aedes echinus* | MW009811*<br>MW009812*<br>MW009813* | - | MW008765* |
| *Aedes geniculatus* | MT808437 | MT808465 | MT993491 |
| *Aedes pulcritarsis* | MT808426 | MT808454 | MT993482 |
| *Aedes rusticus* | MT808432 | MT808460 | MT993495 |
| *Aedes sticticus* | MT808442 | MT808470 | MT993497 |
| *Aedes vexans* | MT808428 | MT808456 | MT993484 |
| *Anopheles algeriensis* | MT808436 | MT808464 | MT993490 |
| *Anopheles claviger* | MT808441 | MT808469 | MT993496 |
| *Anopheles plumbeus* | MW009808*<br>MW009809*<br>MW009810* | - | MW008764* |
| *Anopheles pseudopictus* | MT808433 | MT808461 | MT993487 |
| *Anopheles sacharovi* | MT808434 | MT808462 | MT993488 |
| *Anopheles superpictus* | MT808443 | MT808471 | MT993498 |
| *Anopheles melanoon* | MT808424 | MT808452 | MT993480 |
| *Coquillettidia buxtonii* | MT808430 | MT808458 | MT993486 |
| *Coquillettidia richiardii* | MT808427 | MT808455 | MT993483 |
| *Culex impudicus* | MT808438 | MT808466 | MT993492 |
| *Culex modestus* | MT808439 | MT808467 | MT993493 |
| *Culex perexiguus* | MT808440 | MT808468 | MT993494 |
| *Culex pipiens* | MT808425 | MT808453 | MT993481 |
| *Culex theileri* | MT808435 | MT808463 | MT993489 |
| *Culiseta annulata* | MT808429 | MT808457 | MT993485 |
| *Culiseta longiareolata* | MT808423 | MT808451 | MT993479 |
| *Uranotaenia unguiculata* | MT808444 | MT808472 | MT993499 |

\*partial sequences (Sanger sequencing)

**Supplementary Table 2.**
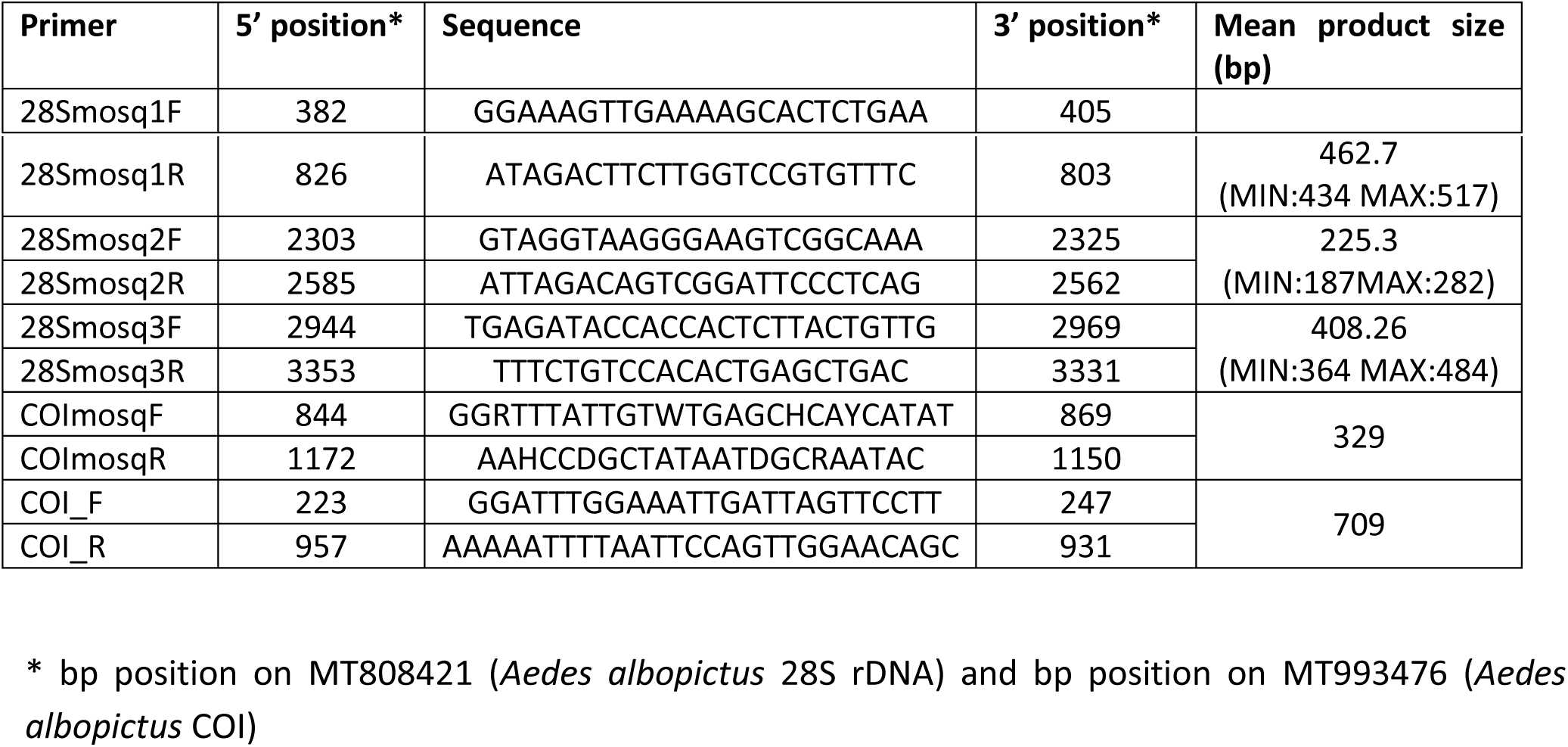
Primers sequences, positions and product sizes.

**Supplementary Table 3.**
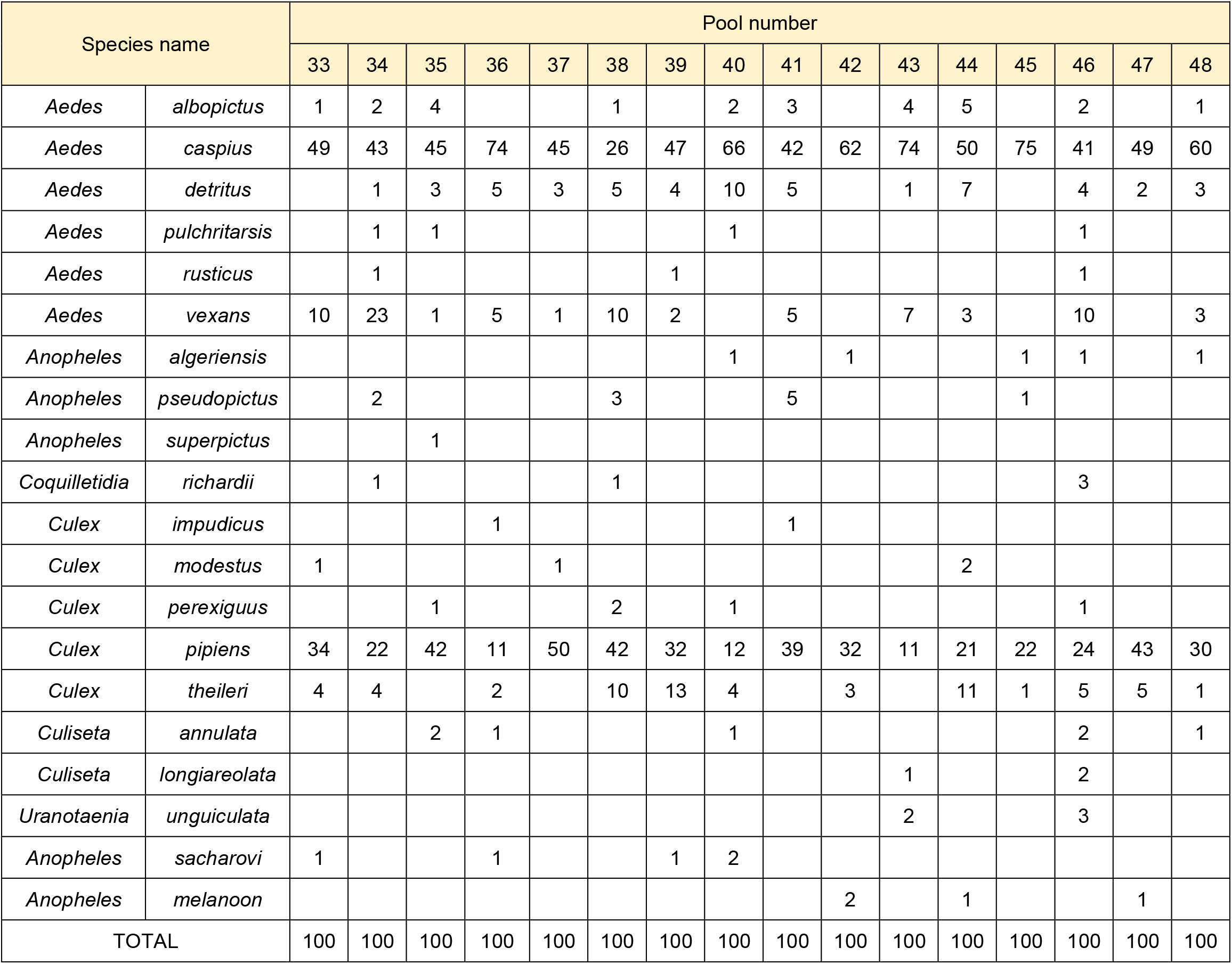
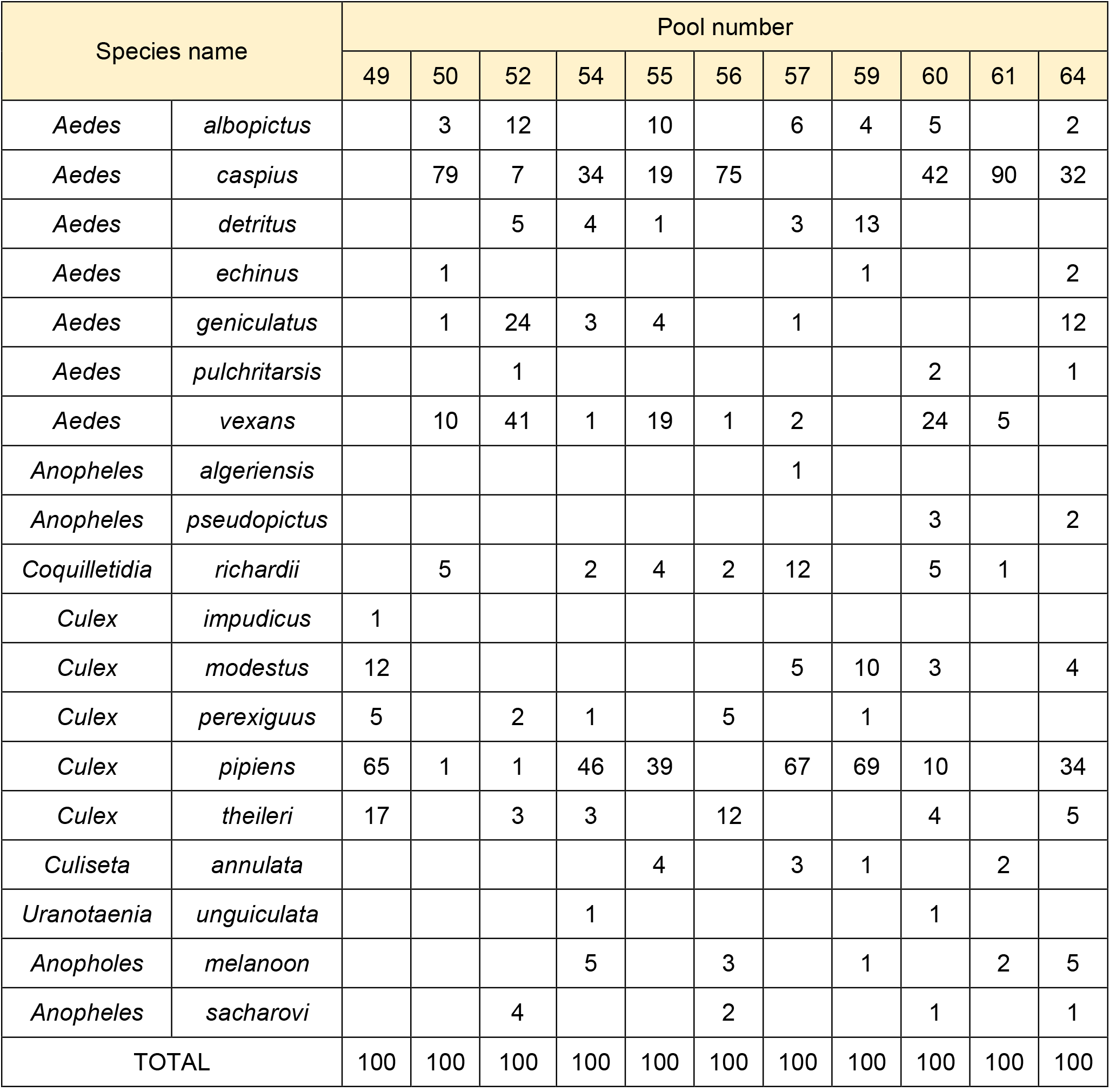
Pools composition according to morphological keys and single-specimen COI identification.

## Supplementary Figures

**Supplementary Figure 1.**
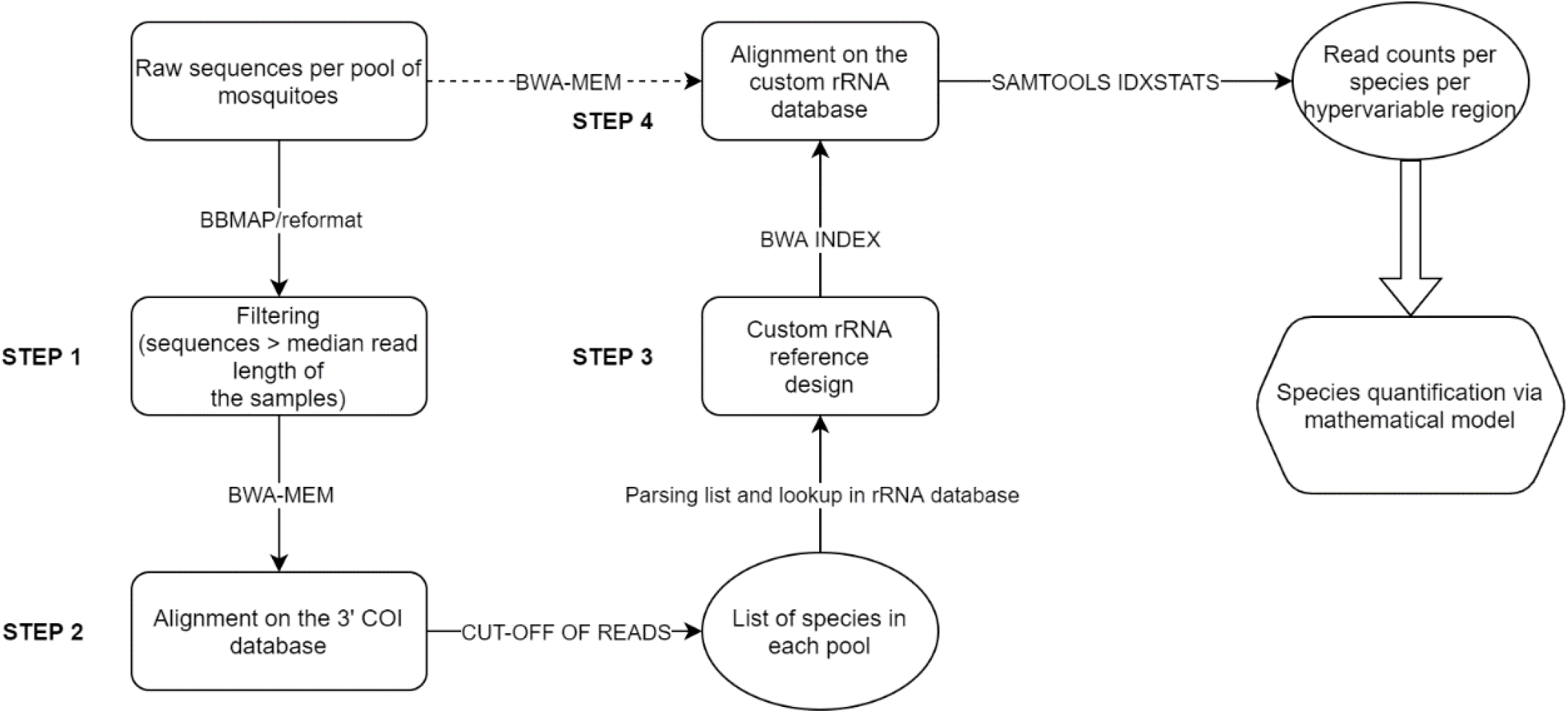
Metabarcoding bioinformatics pipeline logical diagram.

**Supplementary Figure 2.**
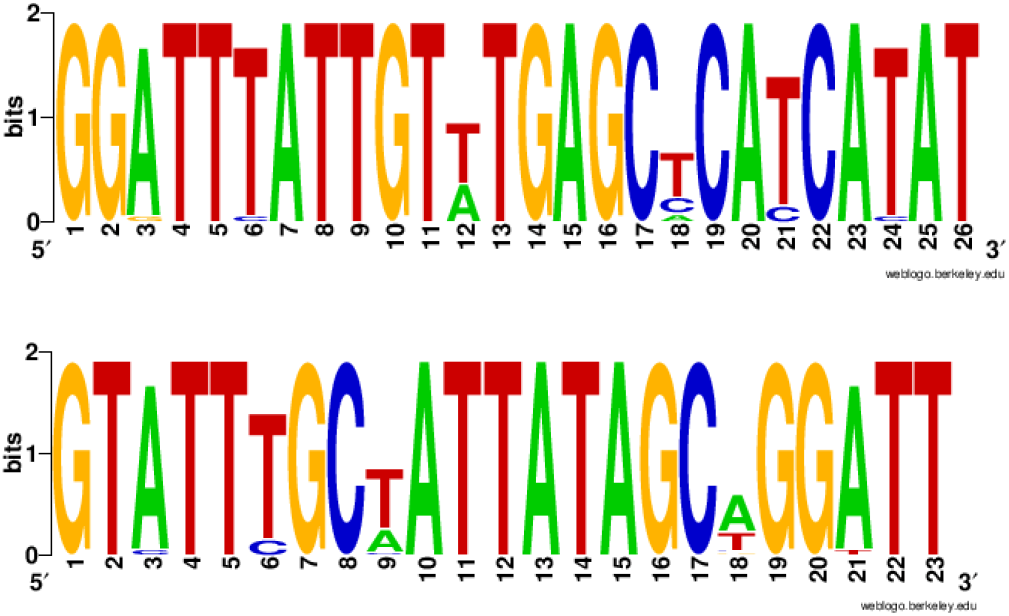
Sequence logos depicting COI primer (COImosqF/COImosqR) annealing sites on the 24 mosquitoes with complete COI CDS. https://weblogo.berkeley.edu/logo.cgi

